# Application of Explainable AI in Neuroscience: Enhancing Autism Screening

**DOI:** 10.64898/2026.02.13.705821

**Authors:** Oana Geman, Sara Sharghilavan, Hadi Abbasi, Roxana Toderean, Octavian Postolache, Alexandra-Stefania Mihai, Matti Karppa

**Affiliations:** Data Science and AI, Computer Science and Engineering, Chalmers University of Technology, University of Gothenburg, Gothenburg, Sweden; Department of Cognitive Neuroscience, University of Tabriz, Tabriz, Iran; Faculty of Electrical and Computer Engineering, University of Tabriz, Tabriz, Iran; Department of Computers, Electronics and Automation, Stefan cel Mare University of Suceava, Suceava, Romania; Instituto de Telecomunicacoes and Iscte, Instituto Universitario de Lisboa, Lisbon, Portugal

**Keywords:** eXplainable AI (XAI), elctroencephalography (EEG), ERP, Autism, Deep Learning, screening, Shapley - Additive exPlanations (SHAP), LIME - Local Interpretable Model-agnostic Explanations

## Abstract

The main challenges in the life of a child with autism are difficulties in communication, behavior, and social interaction. Early diagnosis of this neurodevelopmental disorder improves patient outcomes by enabling more effective, personalized interventions. This diagnosis can sometimes be difficult, especially in very young children. Non-invasive, relatively accessible, and able to reflect neural function in real time, electroencephalography (EEG) shows promise in the detection of Autism spectrum disorders (ASD). However, because EEG data is still difficult for experts to understand, machine learning and artificial intelligence (AI) are beginning to be used in this field as well. In this paper, a ResNet+BiLSTM hybrid deep network was applied and achieved high accuracy in distinguishing individuals with autism from neurotypical subjects. Since AI models typically provide predictions without clear explanations, this study employs explainable AI (XAI) methods such as SHAP (SHapley Additive exPlanations) and LIME (Local Interpretable Model-agnostic Explanations) to clarify their decision-making.Delta, theta, alpha, beta, and gamma waves, as well as ERP components P100, N100, P200, MMN, and P600, were analyzed in the two neurotypical and autistic groups that were compared in this study using EEG recordings. By integrating SHAP and LIME, the system achieved both accurate classification and transparent explanations, pointing to EEG- and ERP-based features as reliable biomarkers for ASD.

## 1 Introduction

Autism spectrum disorder (ASD) is a spectrum of neurodevelopmental disorders [1] characterized by deficits in social interactions and interpersonal communication and restrictions in interests or repetition of activities [2]. Furthermore, individuals with autism spectrum disorder exhibit substantial deficits across multiple cognitive domains [3], with particularly pronounced impairments in perceptual functioning, information-processing mechanisms, and attentional regulation [4]. Although a combination of genetic and environmental factors may contribute to the development of this disorder, its exact cause remains unknown [5]. Studies have shown that the global prevalence of this disorder is approximately 0.6%, which has been increasing over time [6]. Appropriate and early therapeutic intervention in childhood can improve the symptoms of this disorder to an acceptable extent [7].

The increase in the prevalence of this disease has entailed high costs for the recovery and education of these individuals and, at the same time, has generated psychological problems for their families [8]. Thus, new technologies could help doctors and therapists with early diagnosis and choosing the right tools and methods of therapy, improving the quality of life of the children and their families [9]. One of these techniques is artificial intelligence. Artificial intelligence techniques facilitate the analysis of extensive multimodal datasets, offering objective insights into the core mechanisms of brain function. This approach could support earlier and more precise diagnosis of neurological disorders and guide more effective intervention strategies [10].

Explainable AI (XAI) encompasses a set of techniques designed to clarify the decision-making processes of complex models, providing interpretable, post-hoc insights even when the underlying system operates as a non-transparent black box [11, 12]. Unlike inherently interpretable models such as linear regression or decision trees, XAI methods allow researchers and clinicians to understand the rationale behind predictions generated by models such as deep neural networks [13, 14, 15]. By enhancing transparency and interpretability, XAI can increase trust in AI-based tools and support their adoption in clinical and therapeutic contexts, ultimately improving decision-making in the care of individuals with ASD [16].

In the field of neuroscience, explainability is particularly crucial, as medical professionals need to understand which specific elements of neural signals, such as certain frequency bands or connectivity patterns, contributed to the model’s output. Linking these features to known neurobiological processes is essential for both scientific understanding and clinical application [17]. Among the most widely used XAI tools are SHAP (SHapley Additive exPlanations) and LIME (Local Interpretable Model-agnostic Explanations) [18], as well as attention mechanisms in neural networks [19] and models based on symbolic reasoning [20]. These techniques help transform machine learning algorithms into reliable solutions that can support both scientific analyses and clinical decision-making.

Several studies have examined EEG and ERP markers in ASD. EEG analyses have revealed altered power spectra in multiple frequency bands [21, 22], atypical connectivity patterns [23, 24], and abnormal local synchronization in children with ASD compared to neurotypical peers [25, 26]. ERP studies have highlighted differences in components such as P100, N100, P200, MMN, and P600, reflecting alterations in early sensory encoding as well as higher-level cognitive processing [27-30]. Moreover, previous work has explored the use of machine learning and deep learning models for classifying EEG data [31-34], demonstrating promising performance in distinguishing ASD from neurotypical participants [35]. However, many of these models lack interpretability, limiting their clinical applicability. To address this limitation, explainable AI (XAI) methods such as SHAP and LIME have been proposed to provide insight into model decision-making. Despite these advances, integrating EEG and ERP features within a unified, interpretable deep learning framework remains an open challenge.

This study provides a comprehensive examination of the methods and analyses employed. The experimental design and preprocessing pipeline are described in detail, along with a thorough account of how EEG and ERP features were processed. The deep learning framework is also explained, highlighting key aspects of the network architecture and training procedures. A major focus of this work is model interpretability; to this end, SHAP and LIME analyses are combined to provide a more complete understanding of the factors influencing the model’s predictions.

In this study, we build upon the EEG-based classification framework by integrating additional neurophysiological markers and enhancing the learning architecture. Specifically, this study introduces event-related potential (ERP) features alongside EEG signals, focusing on components such as P100, N100, P200, MMN, and P600, which capture both early sensory encoding and higher-level cognitive processes. By combining continuous EEG dynamics with discrete ERP responses, the model leverages complementary sources of information that strengthen its discriminative power and clinical interpretability.

Methodologically, the architecture was improved through a dual-branch framework. The first branch processes raw EEG using a ResNet-BiLSTM backbone augmented with an attention mechanism, enabling the model to capture complex spatio-temporal dependencies more effectively. The second branch is dedicated to ERP-derived features, where both amplitude and latency measures of clinically meaningful components are modeled through a dense neural network. The outputs of these two branches are then fused, producing a unified representation that balances low-level temporal dynamics with high-level neurocognitive markers.

Another key advancement is the use of a dual interpretability strategy. In addition to SHAP, which was employed to analyze EEG channel contributions, the present study incorporates LIME to cross-validate feature relevance and highlight ERP component importance. This multi-layered XAI framework provides not only spatial maps of EEG electrode contributions but also identifies specific ERP components that play a decisive role in distinguishing autistic from neurotypical participants.

## II. Methods

Electroencephalography (EEG), as a continuous and non-invasive technique for recording brain activity [36], provides a valuable tool for investigating neural dynamics and functional abnormalities associated with autism spectrum disorder (ASD) [37].

Several studies in recent years have shown that brain electrical activity, as measured by EEG, differs in people diagnosed with ASD, particularly in how different brain regions collaborate [38, 39]. These differences in EEG coherence show promise as potential biomarkers, but the scientific community has not yet reached a clear consensus on their applicability in clinical practice. A recent example comes from Mayor T. and his collaborators [40], who proposed a method that makes the decisions of neural networks used in EEG analysis for emotion recognition “intelligible.” Their study highlights how these automated systems can identify relevant patterns, which could help in understanding emotional processing in autism. At the same time, other studies have observed a reduction in functional connectivity in the theta and alpha frequency bands in children with ASD, suggesting that these EEG signals may serve as indicators for early detection of the disorder [41].

The human brain functions in a complex balance of electrical frequencies, each with a specific role in cognitive processes. In autism, this “orchestra” seems to function differently. For example, activity in the delta band (0.5–4 Hz), which occurs mainly during deep sleep and is associated with neuronal regeneration, has been reported to be unusually active in some children with ASD even when awake, a possible sign of delayed brain development or weak inhibitory mechanisms [42]. Theta activity (4–8 Hz), related to attention, memory, and emotion regulation, is also increased in certain regions of the brain in these children, especially in the frontal and central areas. This may be related to difficulties in maintaining concentration and adapting quickly to stimuli [43]. The alpha band (8–12 Hz), considered a sign of relaxation and control over external stimuli, also shows changes in children with ASD: either its intensity is reduced, or its distribution across the scalp is different from that of neurotypical children [44]. These changes may reflect difficulties in the internal organization of brain networks and increased sensitivity to the environment [45].

In ASD research, attention has shifted beyond signal amplitude to also include EEG coherence, which reflects the degree of communication between different brain regions. Studies consistently report altered patterns of coherence in individuals with ASD. For example, children with autism often show increased local connectivity (hyperconnectivity) in nearby brain regions, especially during early development [46], while long-range connectivity, such as between the two hemispheres or between frontal and parietal lobes, tends to be weakened (hypoconnectivity) [47]. These imbalances may have a strong clinical relevance: reduced connectivity across distant brain areas has been linked to difficulties in integrating complex social or cognitive information, while excessive local synchronization may lead to rigid thinking patterns and a lack of cognitive flexibility [41].

### A. Data acquisition

This study involved a group of 13 children clinically diagnosed with high-functioning autism, aged between 7 and 12 years (including 2girls and 11 boys, average age 9.7 years). For comparison, we included a control group of 13 neurotypical children, selected to match the age range of the experimental group (including 4 girls and 9 boys, average age 9.3 years). All participants in the control group were right-handed and had no documented neurological or psychiatric disorders. All participants were native Romanian speakers and had a normal hearing threshold of 25 dB or better at frequencies from 250 to 8,000 Hz. Before the experiment began, the entire experimental process and its purpose were explained to the parents or legal guardians, and informed consent was obtained from them following the Declaration of Helsinki [48]. All protocols of this experiment were approved by the ethics committee of Stefan sel Mare University.

All experimental sessions were conducted in a quiet, noise-free, and distraction-free environment. The experimental sessions lasted approximately 10 min and the stimuli consisted of short sentences describing objects in Romanian, manipulated by various acoustic features of distance, tempo, pitch, white noise, and direction. Before the start, subjects were asked to remain still, avoid sudden movements, and actively listen to the auditory stimuli. All recordings were made using a 19-channel OpenBCI headset, configured according to the internationally recognized 10-20 system for electrode placement (Figure 1).

**Fig. 1.**
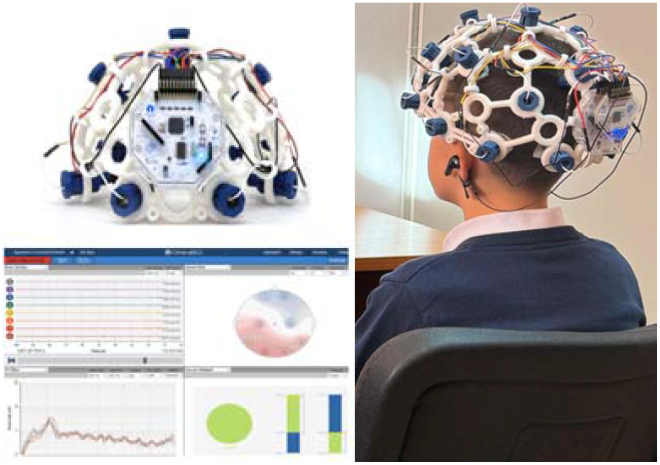
Electrode placement based on the international 10–20 system (19-channel OpenBCI montage). [49]

While auditory stimuli were presented via a Python-based user interface that ensured precise timing, ERP markers were simultaneously embedded in the EEG signal to allow for fine-tuning of neural responses with stimulus onset. EEG data were recorded with a preamplifier bandwidth of 0.1–70 Hz (preserves the EEG bands of interest, removes high-frequency drift and noise), a sampling rate of 250 Hz (sufficient to cover 70 Hz without aliasing, efficient in storage/processing), and a resolution of 24 bits and 140 nanovolts (ensuring fine details of EEG signals at the microvolt level) by the OpenBCI-GUI system and Cyton Board. EEG was recorded using a standard electrode array from positions F3, F4, C3, C4, T3, T4, T5, T6, P3 and P4, according to the international 10-20 system. The ground electrode was placed at Fpz and reference electrodes were placed at A1 and A2. In this study, we analyzed the P100, N100, P200, MMN and P600 components, which reflect early sensory and higher-level cognitive processing in response to speech stimuli in participants [49]. During network training, data from both resting and working states were included to help the model learn patterns that have broader applicability and are not limited to a single condition. EEG signal preprocessing.

### B. EEG signal preprocessing

Before applying any filters, the raw EEG recordings were carefully inspected to identify and remove segments affected by clear artifacts, such as pronounced eye movements, muscle activity, or electrical interference. Participants whose data contained excessive restlessness or persistent, non-correctable artifacts were excluded from further analysis. To reduce noise while preserving the relevant neural signals, we applied a series of filters: a high-pass filter at 1 Hz to eliminate slow baseline drifts, a low-pass filter at 40 Hz to suppress high-frequency activity (including muscle-related noise), and a notch filter at 50 Hz to remove interference from the power line.

Although, in theory, a low-pass filter with a sharp cutoff at 40 Hz could substantially attenuate the 50 Hz component, the roll-off of real filters is never perfectly abrupt. As a result, residual 50 Hz noise may still affect the recordings. Therefore, an additional notch filter was implemented to more effectively suppress line noise. After preprocessing, the cleaned signals were segmented into non-overlapping 2-second epochs, ensuring uniformity and comparability in the subsequent spectral analyses across participants.

To reduce contamination from unwanted sources, a Butterworth band-pass filter between 0.1 and 30 Hz was applied, effectively attenuating low-frequency noise arising from movements and slow drifts, as well as high-frequency disturbances such as electrical interference and muscle activity. This ensured that the frequency range most relevant to neural activity was preserved. In addition, Independent Component Analysis (ICA) was used to decompose the EEG signals into statistically independent components. Since artifacts such as blinks, eye movements, and muscle activity are largely independent of genuine neural activity, this approach enabled their systematic identification and removal.

An automated procedure was employed to detect contaminated epochs by applying amplitude thresholds of ±100 µV together with variance-based criteria, that is, identifying segments where the signal variance exceeded a predefined threshold, indicating abnormal fluctuations likely caused by artifacts. Segments flagged as artifactual were excluded from subsequent analyses.

For each participant, the preprocessed data were then transformed using the fast Fourier transform (FFT), allowing the calculation of power spectra within the conventional frequency bands: delta (0.5–4 Hz), theta (4–8 Hz), alpha (8– 12 Hz), and beta (13–30 Hz) (Figure 2).

**Fig. 2.**
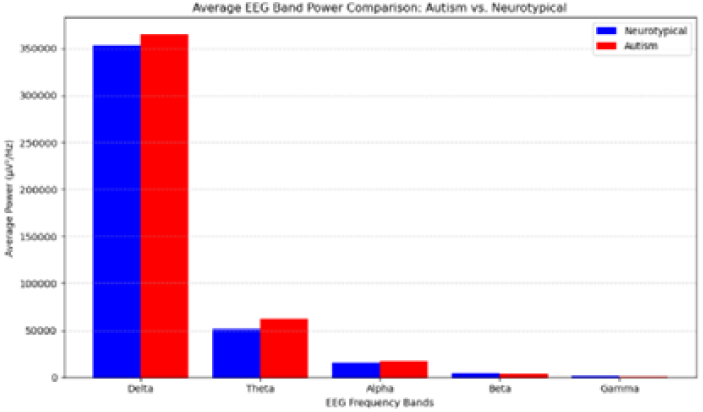
Neurotypical and Autism comparison of average EEG band power across delta, theta, alpha, and beta frequencies.

Beyond spectral analysis, we also examined coherence to capture the degree of functional synchronization between different brain regions, such as frontal–parietal and frontal– temporal connections. These computations were performed for both the experimental and control groups, after which the results were statistically compared. The findings indicated that autism is characterized by reduced coherence across the lower and middle frequency ranges (delta, theta, alpha, and beta), suggesting that the communication between major cortical areas is weaker and less well synchronized (Figures 3–6). A particularly notable reduction at low frequencies (delta and theta) points to limited integration between frontal regions, which support executive control, and parietal regions, which contribute to sensory processing and attentional regulation. In children with autism, this pattern may underlie challenges in sustaining attention and flexibly adapting their responses to external stimuli.

**Fig. 3.**
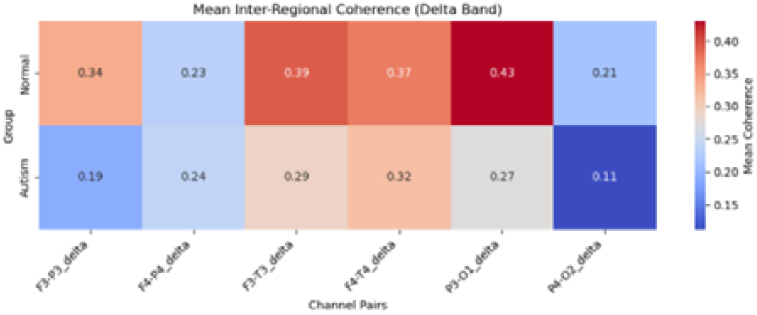
Mean inter-regional coherence in the delta band for ASD and Neurotypical groups [37].

**Fig. 4.**
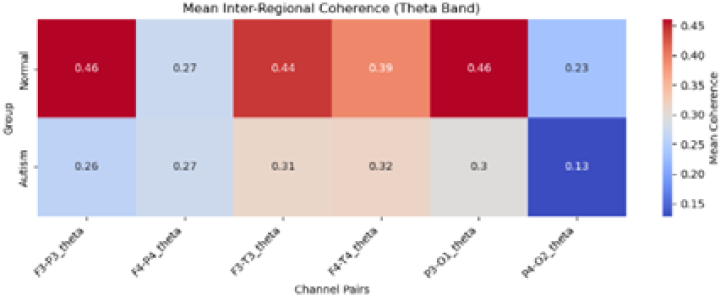
Mean inter-regional coherence in the theta band for ASD and Neurotypical groups [37].

**Fig. 5.**
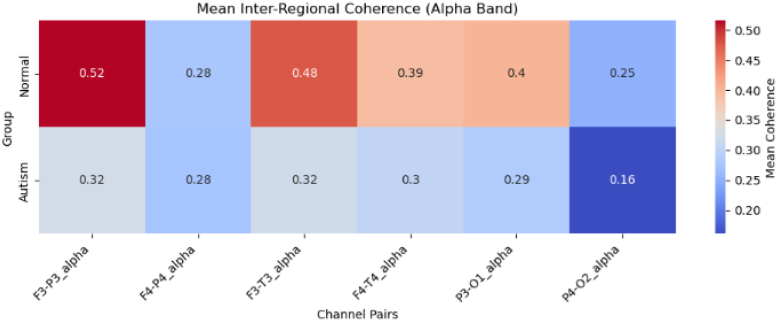
Mean inter-regional coherence in the alpha band for ASD and Neurotypical groups [37]

**Fig. 6.**
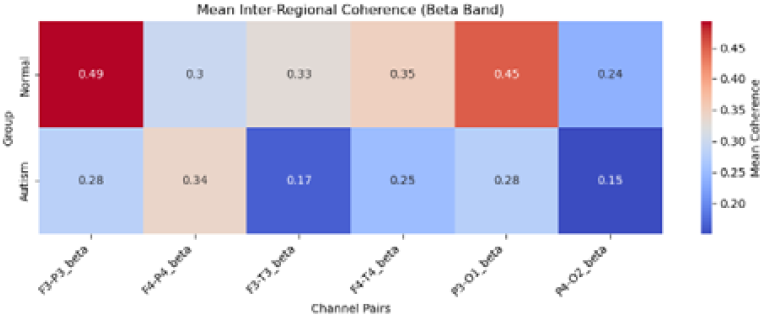
Mean inter-regional coherence in the beta band for ASD and Neurotypical groups [37].

Lower coherence in the theta and alpha bands between frontal and temporal regions has been repeatedly associated with deficits in communication and language, as well as with difficulties in processing faces and social cues, functions that are particularly vulnerable in autism. A reduction in frontal– temporal coherence within the theta range, in particular, appears to be related to challenges in understanding others’ intentions and in generating adequate emotional responses (Figures 3–6).

A reduction in coherence within the alpha and beta bands points to weakened functional connectivity, a finding frequently linked to difficulties in integrating social and cognitive information.

Conversely, in the higher gamma range, localized increases in coherence can sometimes be observed, suggesting excessive synchronization within restricted neural circuits that occurs at the expense of broader network integration. Patterns of altered coherence are being recognized as reliable EEG biomarkers with utility in the early identification of autism and in monitoring clinical response to interventions. This analytic approach makes it possible to localize deficits in network coordination, thereby offering clinicians a framework for tailoring interventions to specific functional domains, including attention, communication, and sensory integration [37].

### C. ERP Components and their Clinical Significance

Event-related potentials (ERPs) highlight how the brain processes sensory and cognitive events over time. Research shows that early and late ERP components are different in people with autism, highlighting atypical patterns of neural functioning [50].

Mismatch negativity (MMN), which occurs approximately 100-250 ms after the presentation of an auditory stimulus and reflects the brain’s automatic detection of changes in auditory flow [51]. Alterations in MMN, as found in autism, highlight that rapid, pre-attentive analysis of auditory input, a crucial skill for language acquisition, may be impaired [52].

P100 occurs between 80 and 130 ms and represents the first stage of sensory recording [53]. In autism, both the amplitude and latency of this component are often altered, reducing neural efficiency in these early stages of processing [54].

P200 is a positive wave that occurs around 150-275 ms and is related to the presentation of auditory or visual stimuli. When listening to read sentences, this wave is stronger when words appear in predictable sentences, especially in the left hemisphere, which plays an important role in anticipatory language processing [55].

N100 occurs between 90 and 150 ms and plays a central role in selective attention, helping the brain to eliminate irrelevant information [56]. In autism, the brain’s lack of discrimination of this component indicates difficulties in filtering distracting stimuli and focusing on more important information [57].

P300, appearing in the 300-600 ms range, stands out as one of the most studied ERP components. It is related to higher cognitive functions such as attention, working memory updating, and stimulus evaluation [58]. In autism, lower amplitudes for P300 are reported, highlighting attention deficits and difficulties in integrating new information into ongoing cognitive activity [59].

Data used for research and diagnosis of ASD is very sensitive by its very nature, presenting multiple ethical challenges for use of AI. Patients’ privacy must be preserved, in addition to ensuring their informed consent, mitigating unfair algorithmic bias, and upholding accountability of the practicioners and providers of AI systems. This makes it crucial to develop transparent and fair AI systems that foster trust of both clinicians and patients’ families, as AI technology is responsibly integrated into healthcare.

### D. ResNET-BiLSTM model

Our method combines knowledge from classical neurophysiology with the analytical power of modern deep learning. Instead of treating these fields as competing alternatives, we integrate them, leveraging the precision of established neuroscience and the adaptability of advanced machine learning to achieve more reliable diagnostic results.

The implementation was done in Python, with Keras and TensorFlow used to build the processing pipeline. Raw EEG data served as input for a ResNet-BiLSTM hybrid model developed to distinguish between autism and typical brain activity.

The ResNet component introduces residual blocks that prevent the vanishing gradient problem, allowing information to bypass intermediate layers. This allows the model to learn both short and transient features of neural activity, as well as broader and longer-lasting patterns within the same signal. BiLSTM extends this capability by analyzing EEG sequences in both temporal directions, an approach that captures complex dependencies over time that are fundamental to understanding brain function [60].

Our study also involves an attention mechanism, which allows the model to highlight time intervals relevant for classification. This is important in autism research, where the most relevant neural responses can be found at specific moments after the presentation of a stimulus.

Another distinctive feature of the system is its dual-input design. On the one hand, it incorporates event-related potentials (ERPs) with well-documented clinical markers such as P100, N100, P200, MMN, and P600. This method processes raw EEG signals, which may also contain patterns that have not yet been identified as biomarkers but may be relevant for diagnosis.

By combining validated neurophysiological markers with unexplored EEG features, the model does more than reproduce existing knowledge, it provides a framework for discovering new diagnostic signatures while strengthening the reliability of established findings.

We used LIME and SHAP for feature interpretation. The proposed EEG classification framework is designed as a dual-branch deep learning architecture that jointly leverages raw EEG time-series and event-related potential (ERP) features to improve discriminative performance. This hybrid approach aims to capture both the fine-grained temporal– spatial dynamics inherent in EEG signals and the neurocognitively interpretable characteristics of ERP components.

In the raw EEG branch, spatial representations are extracted through a series of Residual Network (ResNet) blocks comprising 1D convolutional layers with progressively increasing filter depths (64, 128, 128, 64), each followed by batch normalization and ReLU activation. The residual skip connections mitigate vanishing gradient problems, stabilize training, and enable the network to model more complex hierarchical patterns. The convolutional backbone is followed by stacked Bidirectional LSTM (BiLSTM) layers, which learn long-range dependencies by processing temporal sequences in both forward and backward directions. To further refine the temporal representation, an attention mechanism is integrated to adaptively weight salient time points, ensuring that the network emphasizes segments most relevant to classification.

In the ERP branch, handcrafted features derived from canonical ERP components (P1, N1, P2, MMN, P300, and P600) are employed. For each component, both amplitude and latency measures are extracted, capturing well-established neurophysiological markers that differentiate autistic from neurotypical populations. These features are passed through a fully connected dense subnetwork with ReLU activations, batch normalization, and dropout layers, enabling the model to learn nonlinear relationships among ERP indices while mitigating overfitting.

The outputs of the two branches are concatenated to form a unified latent representation, which is subsequently processed by additional dense layers (128 and 64 neurons) with dropout regularization. A final Softmax layer produces probabilistic predictions for the binary classification task (autism vs. control).

For training, both EEG and ERP inputs are standardized to zero mean and unit variance. Data are split using an 80/20 train–test partition, with an additional validation set drawn from the training data. The model is optimized using the Adam optimizer (learning rate = 0.001) with categorical cross-entropy loss as the objective function. Training is conducted for up to 30 epochs with a batch size of 32, while early stopping (patience = 10) monitors validation loss to prevent overfitting. To further enhance robustness, results are averaged across multiple experimental runs with varying hyper parameters, ensuring reproducibility and stability of reported performance.

This multi-branch design allows the model not only to achieve higher predictive accuracy compared to single-input architectures but also to provide complementary insights: the raw EEG branch captures distributed spatiotemporal patterns, while the ERP branch introduces neurocognitive interpretability aligned with established biomarkers of autism.

Various experiments have been conducted to select the hyperparameters to achieve the best result [57], [37] . The results reported in this study are the average of several experiments.

### E. Feature interpretation using XAI

Bringing explainable AI (XAI) into EEG and ERP analysis for early ASD detection has the potential to improve both scientific understanding and clinical confidence. Tools like SHAP and LIME, which work across different model types, can shed light on why a system makes a particular prediction by showing how features such as spectral power, connectivity patterns, or ERP timing influence the outcome. This kind of transparency not only helps researchers audit complex models and spot misleading patterns, but also points toward new neurophysiological insights that can be tested in future studies. Still, striking the right balance between accuracy and interpretability is not straightforward, especially when working with deep neural networks or time-dependent brain signals.

Moving forward, research should focus on carefully assessing the reliability of explanations, developing methods that capture temporal and layer-specific information in DNNs, addressing feature overlap, and involving clinicians in the loop to ensure that explanations truly support decision-making without sacrificing model performance.

For clinicians, trust in AI systems depends on more than just accuracy, it requires clear reasoning behind each prediction. Neurologists and psychologists, for instance, need to know why a model identifies a child as high-risk, not simply that it does. Methods like SHAP and LIME address this by providing case-specific explanations that highlight which EEG or ERP features, such as theta power, P300 latency, or connectivity patterns, most influenced the decision.

## III. Results

EEG analysis was conducted on a cohort of 13 children diagnosed with ASD and a matched control group of 13 neurotypical children, balanced for age and gender. Following filtering and preprocessing, the EEG data were analyzed using deep learning models, and interpretability was examined through XAI methods, including LIME and SHAP. After model training, classification performance was evaluated on the test set, with accuracy serving as the primary performance metric. The model achieved a classification accuracy of **0.9627** on the test set.

Figure 7 presents the confusion matrix of classification results on the test set. The model demonstrated excellent performance, with both high overall accuracy and balanced precision and recall. Only a small number of misclassifications were observed (11 false positives and 8 false negatives). We prevented the overfitting issue by ensuring that data from the same individuals only appeared in either the training set or the test set, but never in both. Notably, the classifier exhibited slightly higher recall in detecting ASD cases, while also maintaining strong performance in identifying neurotypical controls. This balance underscores the reliability of the model in distinguishing between the two groups.

**Fig. 7.**
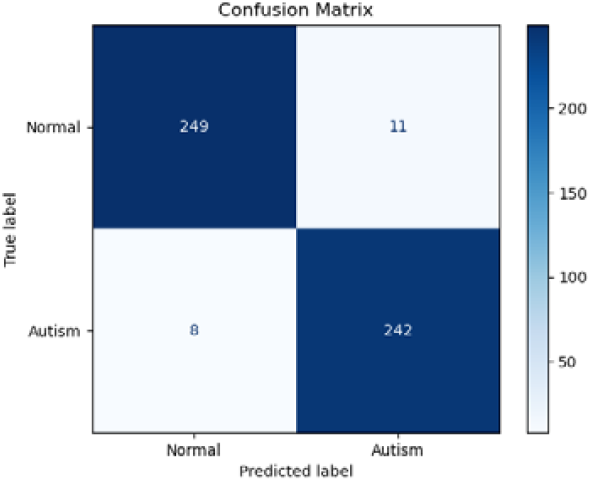
Confusion matrix showing the classification performance on the test set

Key metrics from this confusion matrix:

Accuracy = (249 + 242) / (249 + 11 + 8 + 242) = 491 / 510 ≈ 96.27%

Precision (for Autism) = 242 / (242 + 11) ≈ 95.65%

Recall (for Autism) = 242 / (242 + 8) ≈ 96.80%

F1-Score (for Autism) = 2 × (Precision × Recall) / (Precision + Recall) ≈ 96.22%

To gain further insight into the model’s decision-making process, XAI techniques such as SHAP were employed to interpret the contribution of manually extracted ERP features to classification outcomes. After training the convolutional neural network to categorize participants into ASD versus neurotypical groups, SHAP values were computed to quantify feature importance. SHAP provides an estimate of how much each feature contributes to the final prediction, based on cooperative game theory. For each participant, the system was able to explain the specific factors influencing their classification, with electrode-level scores indicating the degree to which each channel pushed the decision toward ASD or toward the neurotypical class. Positive values reflected contributions to ASD classification, while negative values contributed to the control group.

These feature contributions were visualized topographically across the scalp to identify the spatial distribution of relevant neural activity. Such maps revealed the “spatial signature” of ASD within EEG patterns. Features that consistently emerged as important across both SHAP and LIME were considered robust markers, reducing the likelihood of false interpretations. The resulting topographic maps provided clear visual cues regarding the brain regions most involved in classification.

In addition to EEG features, further interpretability analyses were applied to models based on fully connected neural networks and LSTMs using SHAP values. These values were calculated for spectral power features across channels and frequency bands, highlighting both positive and negative contributions to classification (Fig. 8a and 8b). As shown in Figure 8a, red areas, primarily over lateral and extend partly into frontal scalp regions, contributed positively to identifying the neurotypical class, whereas blue areas, particularly central and frontal regions, indicated stronger associations with ASD. These spatial patterns provided valuable insights into the regional importance of EEG activity in the model’s decisions. At the group level, SHAP-based visualizations revealed consistent patterns that aligned with existing literature, especially alterations in theta and alpha band activity and evidence of fronto-parietal hypoconnectivity in ASD (Figure 8b).

**Fig. 8.**
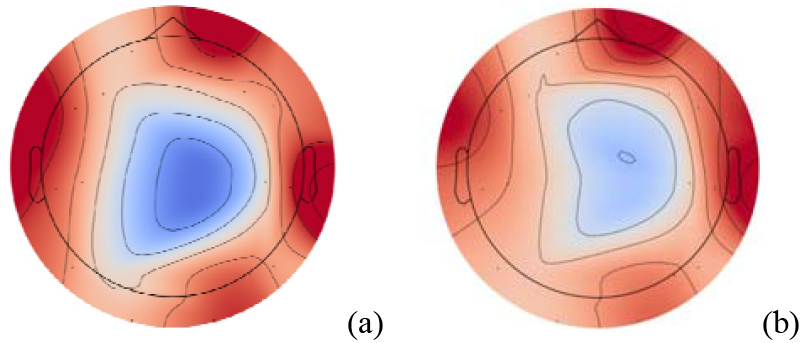
SHAP-based feature importance visualization for EEG inputs. Warmer colors indicate stronger positive contributions to classification. for Neurotypical class (a); and for Autism class (b)

Figure 9 shows the importance of SHAP-based characteristics of ERP components for the two groups (neurotypical vs. ASD). The results highlight that early ERP components, such as P100 and N100, have a relatively greater importance in distinguishing between groups, with stronger contributions observed in the neurotypical class. In contrast, later components, such as P300 and P600, show increased importance in the autism group, suggesting that To avoid overfitting, we need to ensure that the same individuals do not appear in both the training and the test setsatypicalities in higher-order cognitive processing also play a role in classification. These differences illustrate how both early sensory responses and later cognitive components contribute complementarily to the model’s decision-making.

**Fig. 9.**
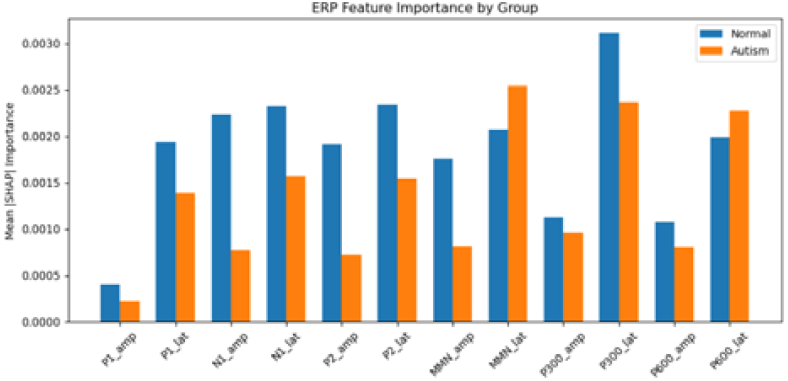
SHAP-based ERP feature importance by group visualization for EEG inputs

To complement SHAP, LIME (Local Interpretable Model-agnostic Explanations) was applied, providing an alternative perspective through localized perturbation of input features. This comparison ensured that feature importance findings were not dependent on a single interpretability method. Figure 10 displays LIME-based visualizations of EEG feature importance, again separated by class.

**Fig. 10.**
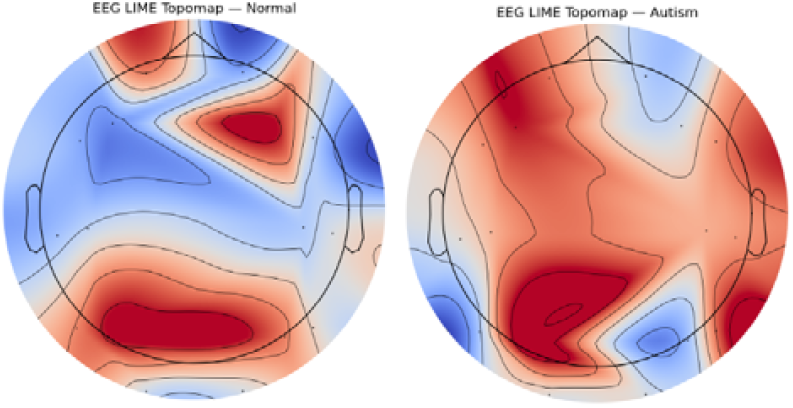
LIME-based feature importance visualization for EEG inputs. Warmer colors indicate stronger positive contributions to classification. for Neurotypical class (a); and for Autism class (b)

LIME-based topographic maps show the importance of EEG features for the two groups. In the neurotypical group (left panel), the red regions are concentrated in the posterior and parietal areas, indicating that these areas contribute more to the classification of subjects as neurotypical. In contrast, in the autism group (right panel), positive contributions are evident in the frontal and temporal regions, suggesting that differences in neural activity in these areas played a more important role in identifying cases of ASD.

For ERP-derived features, both SHAP and LIME were employed to evaluate the relative contribution of different ERP components. Figure 11 presents ERP feature importance by group identified using LIME weights, while Figure 12 and Figure 13 illustrate ERP importance identified through SHAP and LIME, respectively. In these visualizations, blue bars correspond to the neurotypical class and red bars to the ASD class, sorted by relevance to ASD classification. Red stars (⍰) highlight early ERP components, which consistently emerged as the most predictive features across both interpretability methods. The convergence of SHAP and LIME on these components underscores their robustness as candidate early biomarkers of ASD.

**Fig. 11.**
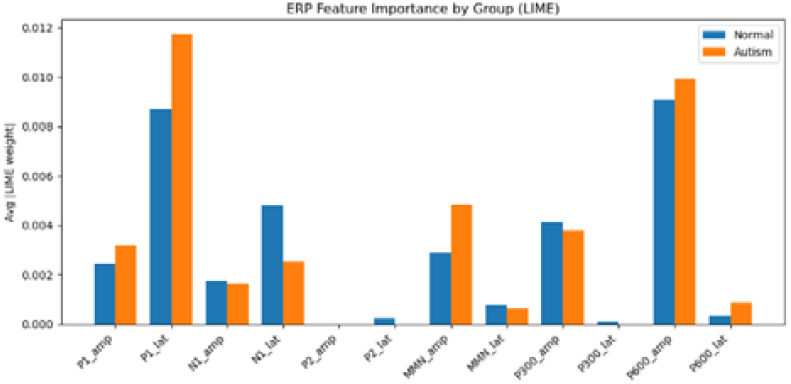
ERP feature importance by group identified using LIME wieght.

**Fig. 12.**
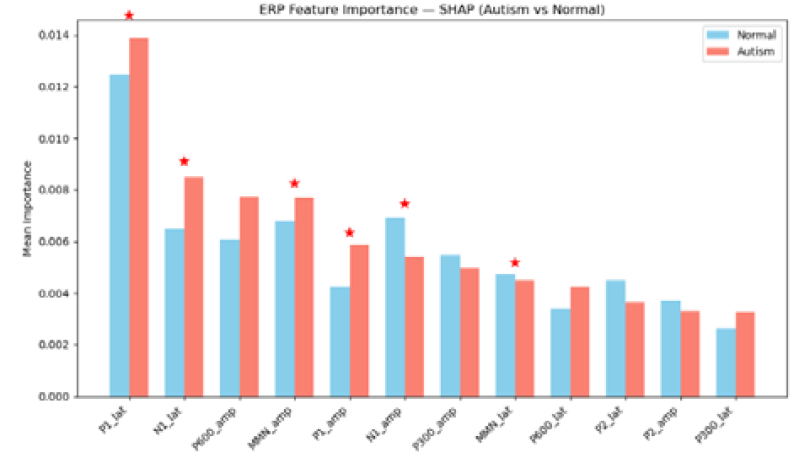
ERP feature importance identified using SHAP.

**Fig. 13.**
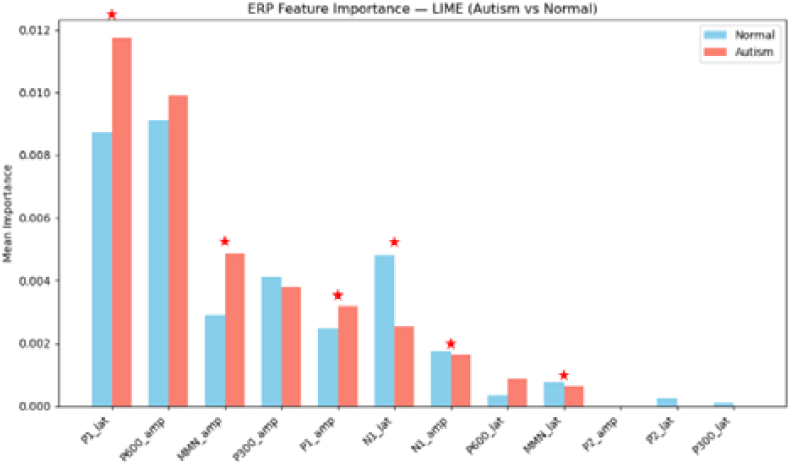
ERP feature importance identified using LIME.

The system automatically identifies the most important ones: brain regions (via topographic maps), ERP components (via importance analysis) and temporal moments (via attention mechanism).

Together, these results provide converging evidence that early sensory-related ERP components (P100, N100, MMN), along with alterations in theta and alpha oscillatory activity and fronto-parietal connectivity, represent important neural signatures of autism. Importantly, the system not only achieved strong classification accuracy but also delivered interpretable predictions at both the group and individual level.

Results can be compared with existing literature to validate whether the model identifies known patterns (e.g. P300 reductions) and to discover potential novel biomarkers [61]. The system provides the explicitable predictions for each patient, a diagnostic confidence through consensus of multiple methods and a guidance for clinicians on the most relevant features.

## IV. Conclusion

Advanced AI models, such as Support Vector Machines, Deep Neural Networks, and Random Forests, have already shown strong performance in distinguishing children with ASD from typically developing peers. Just as importantly, some of these models are being designed to explain their reasoning, which helps clinicians trust and apply AI findings in real-world practice. Aligning AI predictions with established diagnostic criteria ensures these tools complement, rather than replace, clinical expertise.

Recent studies [62], [63] have shown that individuals diagnosed with autism spectrum disorder (ASD) often present distinct patterns of brain activity, especially in the delta (0.5–4 Hz), theta (4–8 Hz), and alpha (8–12 Hz) frequency bands [30]. These differences are not limited to signal intensity; they also involve how various areas of the brain communicate and synchronize with each other. Altered coherence between regions, either reduced connectivity across long distances or excessive local synchronization, has been widely documented.

A major challenge in applying artificial intelligence to clinical neurophysiology lies in the opacity of most models. While many AI systems achieve high accuracy, their internal workings often remain difficult to interpret. This becomes a serious limitation in clinical practice, where understanding why a model made a decision is just as important as the decision itself. To bridge this gap, explainable artificial intelligence (XAI) methods have been developed. These tools allow clinicians to trace the reasoning behind model predictions by identifying which features of the EEG signals were most influential. In our research, we employed SHAP (Shapley Additive Explanations) to interpret predictions from deep learning models, including fully connected networks and LSTMs. This helped us better understand how specific neural features contribute to ASD classification.

Our results indicate that early ERP components such as P100, N100, and MMN, linked to initial sensory and auditory processing, play a key role. Moreover, children with autism in our study showed increased delta and theta activity in frontal areas, a drop in alpha power, and lower functional connectivity between frontal and parietal regions. These patterns may reflect difficulties in attention regulation, information processing, and sensory integration.

By combining EEG analysis with interpretable AI models, we offer a tool that not only supports early identification of ASD but also enhances clinicians’ confidence in using automated systems. Ultimately, this approach can contribute to more tailored and effective intervention plans, adapted to each child’s unique neural profile.

Moreover, by combining ERP-derived features with raw EEG data in a dual-branch deep learning model, and further refining this with an attention mechanism, we were able to build a framework that captures both well-known and potentially new biomarkers with greater reliability. Using two complementary interpretability methods, SHAP and LIME, added an extra layer of confidence to our findings, since the features highlighted as important were confirmed independently rather than relying on a single technique. Overall, this integrated approach not only reached high predictive accuracy but also offered insights that are directly relevant for clinical practice, making a step forward in connecting advanced AI methods with the day-to-day needs of diagnostic support. Autism is complex, and no single type of data can capture its full scope. Future research should focus on combining multiple sources, like brain imaging, behavior, and genetics, to improve diagnostic accuracy. Techniques that merge these data streams can provide richer insights and more robust predictions and early ASD identification.

## Ethical Consideration

All procedures involving human participants were conducted in accordance with the ethical standards of the institutional and/or national research committee and with the 1964 Helsinki Declaration and its later amendments or comparable ethical standards. The study was approved by the Research Ethics Committees of the University of Stefan cel Mare Romania .Written informed consent was obtained from all individual participants. For minors, consent was obtained from their parents or legal guardians.

